# The genetic diversity and populational specificity of the human gut virome at single nucleotide resolution

**DOI:** 10.1101/2024.12.30.630844

**Authors:** Xiuchao Wang, Quanbin Dong, Pan Huang, Shuai Yang, Mengke Gao, Chengcheng Zhang, Chuan Zhang, Youpeng Deng, Zijing Huang, Beining Ma, Yuwen Jiao, Yan Zhou, Tingting Wu, Huayiyang Zou, Jing Shi, Yanhui Sheng, Yifeng Wang, CGVR Consortium, Liming Tang, Shixian Hu, Yi Duan, Wei Sun, Wei Chen, Qixiao Zhai, Xiangqing Kong, Lianmin Chen

## Abstract

Large-scale characterization of gut viral genomes provides strain-resolved insights into host-microbe interactions. However, existing viral genomes are mainly derived from Western populations, limiting our understanding of global gut viral diversity and functional variations necessary for personalized medicine and addressing regional health disparities.

Here, we introduce the Chinese Gut Viral Reference (CGVR) set, consisting of 120,568 viral genomes from 3,234 deeply sequenced fecal samples collected nationwide, covering 72,751 viral species-level clusters (VSCs), nearly 90% of which are absent from current databases. Analysis of single nucleotide variations (SNVs) in 233 globally prevalent VSCs revealed that 18.9% showed significant genetic stratification between Chinese and non-Chinese populations, potentially linked to bacterial infection susceptibility. The predicted bacterial hosts of population-stratified viruses exhibit distinct genetic components associated with health-related functions, including multidrug resistance. Additionally, viral strain diversity at the SNV level correlated with human phenotypic traits, such as age and gastrointestinal issues like constipation. Our analysis also indicates that the human gut bacteriome is specifically shaped by the virome, which mediates associations with human phenotypic traits.

This foundational resource underscores the unique genetic makeup of the gut virome across populations and emphasizes the importance of recognizing gut viral genetic heterogeneity for deeper insights into regional health implications.

## INTRODUCTION

The human gut virome is a critical part of the microbiome, regulating bacterial communities and impacting human health[1–5]. Understanding viral genomic diversity and functional variations across global populations may improve viral discovery and treatment, particularly in addressing regional health disparities. Comprehensive reference viral genomes are vital for genome-resolved metagenomics. Recent large-scale metagenomic assemblies from diverse populations have generated valuable viral reference genomes[6–8], enabling deeper insights into strain-level viral diversity and its health implications[9].

Nevertheless, several key aspects remain underexplored, hindering a full understanding of global gut viral genomic diversity and functional variations. First, most available gut viral genomes come from Western populations. While the Unified Human Gut Virome (UHGV) catalog integrates tens of thousands of viral genomes from 12 existing virome datasets[6, 9–19], around 83% of its metagenomic samples are from Western populations, with only about 17% from non-Western sources. Although some studies have explored the Chinese gut virome[7, 20, 21], large-scale genomic assembly are limited. Given that China accounts for nearly 20% of the global population, expanding gut viral genome recovery from Chinese populations is crucial to enhance our understanding of the gut microbiome and increase population diversity in viral databases. Second, most gut virome phylogenetic research focuses on a few viruses with high morbidity[22] or the most abundant in the gut[23]. A comprehensive comparison of viral genomic diversity across populations and their physiological interactions with hosts remains unexplored. Finally, while advances have been made in understanding viral-bacterial interactions[24, 25], the complex interplay between gut viruses and bacteria, and their effects on human phenotypes, are still poorly understood. Here, we present the Chinese Gut Viral Reference (CGVR), a comprehensive catalog of 120,568 viral genomes derived from a nationwide collection of 3,234 fecal samples across mainland China. Comparative analysis with the UHGV revealed that viral genomes from 89.7% of viral species-level clusters (VSCs) in CGVR are unique to Chinese populations, while 4,406 VSCs are shared between Chinese and non-Chinese populations. Phylogenetic analysis of these shared VSCs at single-nucleotide resolution uncovered population-specific genomic variations related to bacterial host infection, with predicted bacterial hosts of dominant viruses displaying distinct genetic features relevant for human health, including multidrug resistance. Furthermore, viral genetic diversity within populations was associated with host physiological traits, largely independent of viral abundance. We also observed that individual-specific gut virome significantly shape interindividual bacteriome variations. Lastly, medication analysis showed that associations between gut viral abundance and human phenotypic traits are mediated by the bacteriome.

## RESULTS

### A metagenomic compendium of 120,568 gut viral genomes from Chinese populations extends global viral diversity

The gut virome is crucial for human health and disease. To enable strain-resolved metagenomics across diverse populations, a comprehensive reference of viral genomes is essential. The Unified Human Gut Virome Catalog (UHGV) recently integrated 12 existing virome catalogs[6, 9–19], establishing itself as the most extensive viral resource to date. However, fewer than 20% of these viral genomes are derived from non-Western populations, which constitute 75% of the global population. This limitation hinders our understanding of global gut viral diversity and its implications for regional health disparities. In this study, we performed gut viral genome assembly using deeply sequenced metagenomic samples (averaging 20 GB) from 3,234 individuals across 30 provinces in China[26].

Following a stringent viral detection pipeline (**Fig. 1a**), we identified 5,531,572 putative viral bins. Quality assessment using CheckV[27] yielded 120,568 medium-to high-quality viral genomes, categorized as follows: 8,667 complete genomes (100% completeness), 55,710 high-quality genomes (>90% completeness), and 56,191 medium-quality genomes (50–90% completeness) (**Fig. 1b,c; Table S1**). For subsequent analysis, we focused on the 64,377 genomes with >90% completeness to address limitations associated with small genome fragments[28] and adhere to UHGV quality standards. The median size of the assembled viral genomes was 82 kb (interquartile range: 43–173 kb) (**Table S1**). We designate this collection of viral genomes as the Chinese Gut Viral Reference (CGVR).

**Figure. 1.**
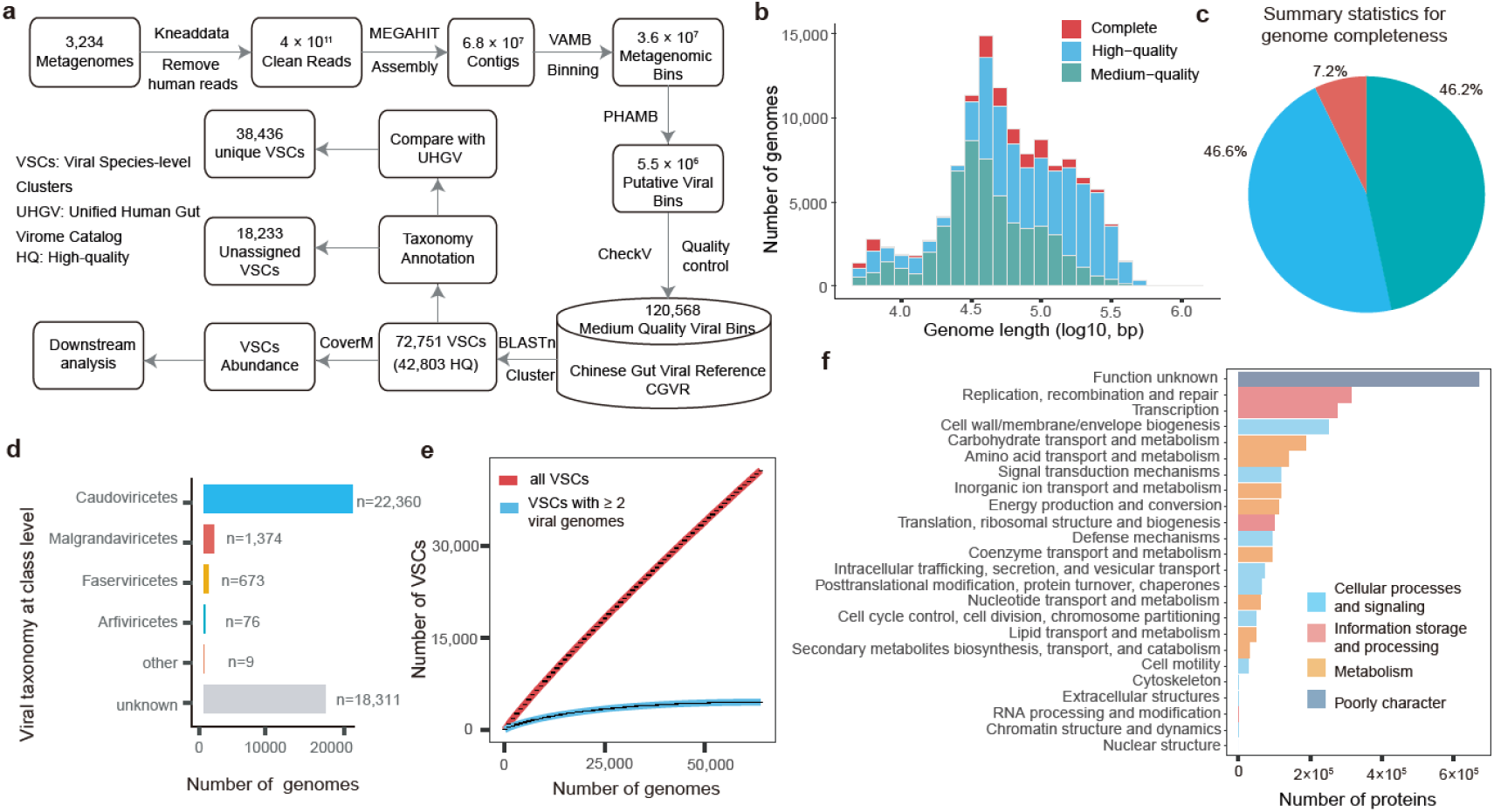
The Chinese Gut Viral Reference (CGVR) extends global viral diversity. **a.** Overview of the CGVR database construction pipeline. A total of 120,568 qualified viral bins were generated, resulting in 72,751 viral species-level clusters (VSCs), including 42,803 high-quality VSCs. **b.** Distribution of assembled viral genomes based on genomic length and quality. **c.** Summary of viral genomes categorized by completeness (complete, n = 8,667; >90% complete, n = 55,710; 50–90% complete, n = 56,191). **d.** ICTV taxonomic annotation of viral genomes in the CGVR, with numbers indicating the total genomes within each viral class. **e.** Accumulation curves for VSCs in the CGVR. Red curves include all VSCs, while blue curves focus on VSCs (n = 7,441) with at least two conspecific viral sequences. **f.** Functional characterization of novel VSCs compared with the UHGV catalog, with bars color-coded by broad COG categories: blue for cellular processes and signaling, red for information storage and processing, orange for metabolism, and grey for poorly characterized.

To classify the taxonomy of our CGVR catalog, we clustered all genomes at 95% average nucleotide identity (ANI) and 85% alignment fraction (AF)[28], resulting in 42,803 viral species-level sequence clusters (VSCs) (**Table S2**). Taxonomic annotation using the ICTV classification tool[29] revealed that 57.22% (24,492) of these VSCs could be classified at the class level, with *Caudoviricetes* (22,360), *Malgrandaviricetes* (1,374), *Faserviricetes* (673), and *Arfiviricetes* (76) being the most prevalent (**Fig. 1d; Table S3**), consistent with recent studies[7, 10, 30]. Notably, 18,311 of the 42,803 (42.78%) VSCs could not be classified into any known viral classes, highlighting the vast, underexplored diversity of gut viruses in Chinese populations. Additionally, rarefaction analysis indicated that the number of VSCs detected has not yet reached saturation, suggesting the presence of additional undiscovered viruses (**Fig. 1e**). However, these are likely to be rarer members of the Chinese gut virome, as the curve approaches saturation when considering only those with at least two conspecific sequences (**Fig. 1e**).

Finally, we examined the functional potential of the 18,311 unclassified VSCs by annotating their protein-coding genes using Prodigal[31] and eggNOG-mapper[32]. A total of 3,531,752 viral proteins were predicted, with 56.79 % (2,005,769) showing homologous genes in the eggNOG database[33], enabling functional decoding (**Fig. 1f**; **Table S4**). Common virus-related proteins involved in processes such as DNA replication, homologous recombination, and transcriptional regulation were frequently identified (**Fig. 1f**). However, 43.21% of protein sequences lacked homology to known sequences in existing databases, highlighting a significant knowledge gap in our understanding of the human gut virome. Overall, the CGVR catalog significantly enhances our understanding of global gut virome diversity, particularly in Chinese populations, and provides a robust foundation for future comparative studies, yielding valuable insights into the potential role of gut viruses in health, disease, and regional disparities.

### Phylogenetic comparison between CGVR and UHGV revealed population-specific gut viruses

We then sought to assess the uniqueness of VSCs in our CGVR compared to non-Chinese viral genomes in the UHGV. To do this, we clustered our 64,377 high-quality viral genomes with 208,604 viral genomes from the UHGV (released in March 2024, uhgv-db-v0.4) using 95% ANI and 85% AF. This resulted in a total of 101,562 VSCs, with only 4,406 (4.34%) overlapping between the CGVR and UHGV (**Fig. 2a**). This finding highlights significant genetic differences in gut viruses between Chinese and non-Chinese populations.

**Figure. 2.**
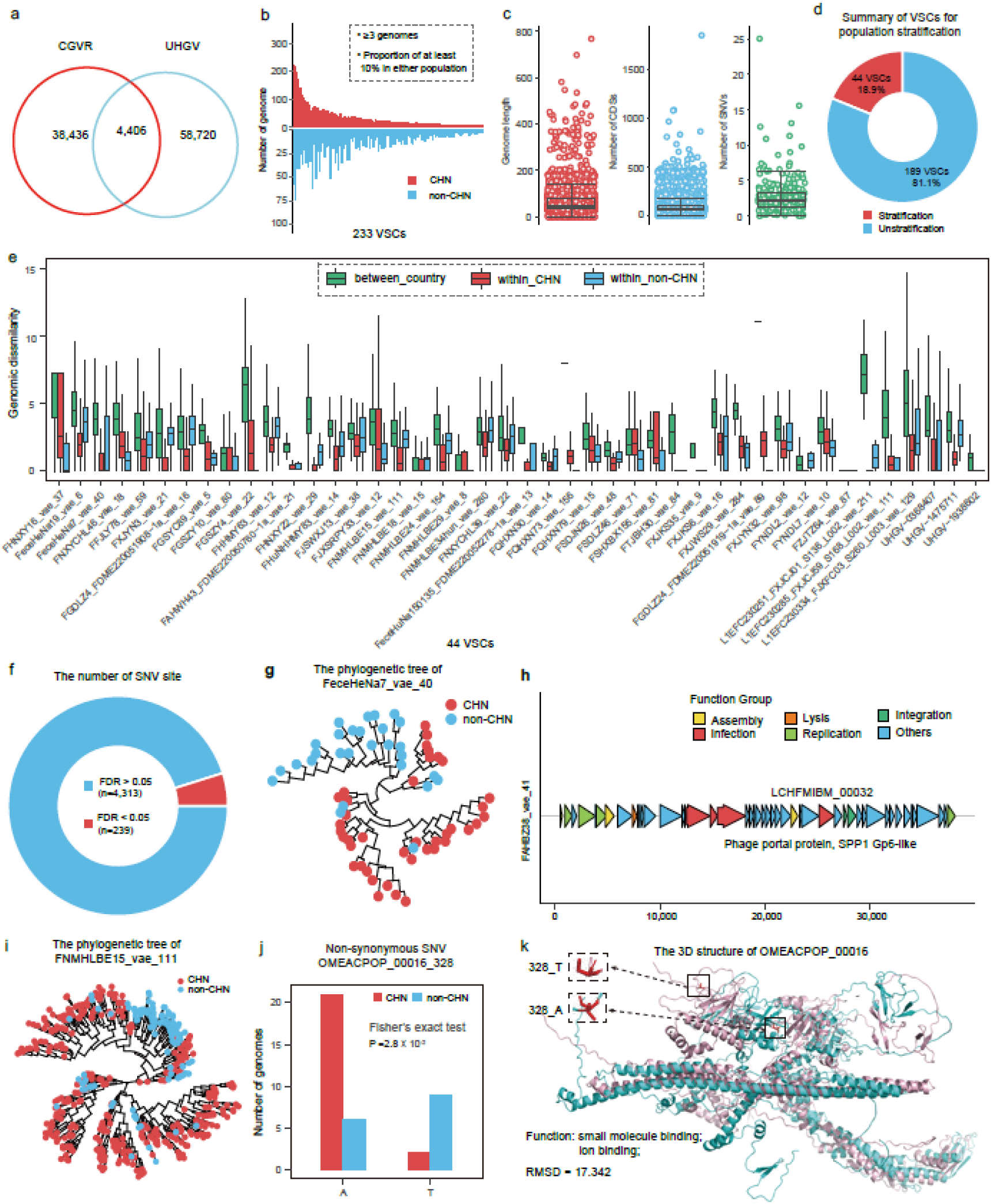
Population-specific viruses and genomic stratification of predominant viruses linked to bacterial host infection. **a.** Overlap of VSCs between the CGVR and UHGV based on 95% ANI and 85% AF. **b.** Summary of viral genomes in 233 commonly presented VSCs across Chinese and non-Chinese populations, defined as having at least three genomes in a given VSC and a minimum of 10% representation in either population. **c.** Characteristics of the 233 commonly presented VSCs. Red dot plot indicates viral genome length, blue dot plot represents the number of viral CDSs, and green dot plot shows detected SNV sites. **d.** Summary of VSCs exhibiting significant genomic stratification between Chinese and non-Chinese populations at single nucleotide resolution. The circular chart shows red for VSCs with significant population stratification (44 VSCs, FDR < 0.05) and blue for those without. **e.** Overview of 44 VSCs with significant genetic stratifications between populations, where green indicates genetic distance between Chinese and non-Chinese populations, red denotes genetic distance within the Chinese population, and blue indicates genetic distance within non-Chinese populations. **f.** Summary of SNVs showing significant enrichment between populations. A total of 4,950 loci from 44 VSCs were analyzed with Fisher’s exact tests, identifying 239 loci that were differentially enriched in either population at FDR < 0.05. **g.** Phylogenetic tree of *FeceHeNa7_vae_40* based on SNVs, with red representing viral genomes from Chinese populations and blue from non-Chinese populations. **h.** Genomic annotation of *FeceHeNa7_vae_40*, highlighting differential SNVs in genes encoding a phage portal protein. Annotated protein-coding genes (arrows) are color-coded by function, including assembly, lysis, integration, replication, and infection. **i.** Phylogenetic tree of *FNMHLBE15_vae_111* based on SNVs, with red indicating genomes from Chinese populations and blue indicating non-Chinese populations. **j.** Summary of viral genomes of *FNMHLBE15_vae_111* with differential allele types in the *OMEACPOP_00016* gene, marked in red for Chinese and blue for non-Chinese genomes. **k.** Predicted protein structure encoded by the *OMEACPOP_00016* gene, with differential physicochemical properties indicated by gray dashed lines.

Notably, 38,436 VSCs from our study, comprising 44,715 viral genomes, did not cluster with any sequences from the UHGV. This substantial divergence underscores the unique viral diversity present in Chinese populations. Given the highly individualized nature of gut viruses at the strain level, developing person-specific viral genome databases is essential for accurate strain tracking[34, 35]. The fact that 89.7% of our VSCs lack close relatives in the UHGV indicates that existing viral catalogs are insufficient for analyzing gut viral strains in Chinese populations. These findings emphasize the necessity of population-specific viral genome databases to capture unique viral diversity, enabling more precise investigations into the gut virome’s role in human health and disease.

### Genomic variations in predominant viruses between populations linked to bacterial host infection

In addition to the VSCs specific to Chinese populations, we examined the 4,406 shared VSCs (**Fig. 2a**) between Chinese and non-Chinese populations for genomic and functional variations. After excluding viral genomes of unknown ethnic origin, we focused on 233 VSCs with at least three genomes and a minimum representation of 10% in either population (**Fig. 2b**). These VSCs averaged 66 kb in genome length and contained around 100 CDSs. Single nucleotide variation (SNV) analysis using Snippy revealed an average of 2,840 SNVs per VSC (**Fig. 2c**).

We then compared within– and between-population genomic dissimilarity using the Kimura 2-parameter model based on SNVs. Among the 233 VSCs, 44 (18.88%) displayed significant population stratification (Kruskal-Wallis test, FDR < 0.05), indicating potential distinct evolutionary trajectories for Chinese and non-Chinese populations (**Fig. 2d, e; Table S5; Fig. S1**).

To assess the functional implications of these genetic differences, we conducted enrichment test on biallelic SNVs (N=4,950), identifying 239 SNVs with significant population differences (FDR < 0.05, **Table S6**), including 181 located within CDSs (**Fig. 2f**). Functional annotation of these genes encompasses various functionalities related to viral lysis, infection and integration (**Table S6, Fig. S2**).

For instance, *FeceHeNa7_vae_40*, a member of *Caudoviricetes*, exhibited two distinct clusters between populations (Fisher’s exact test, P = 6.6×10⁻¹³) (**Fig. 2g**), with eight significant SNVs detected, three of which were within the *LCHFMIBM_00032* gene encoding a phage portal protein (SPP1 Gp6-like) (**Fig. 2h**), crucial for phage infection[36].

Another VSC, *FNMHLBE15_vae_111* from the *Zeta-crassviridae* family, also showed significant genetic stratification (Fisher’s exact test, P = 3.8×10⁻¹⁴) driven by 31 SNVs (FDR < 0.05) (**Fig. 2i**). Notably, a non-synonymous SNV in the *OMEACPOP_00016* gene was enriched in distinct populations, with position 328 showing A in Chinese and T in non-Chinese genomes (**Fig. 2j**). Although this gene lacks functional annotation, predictions using AlphaFold3[37] and DeepFRI[38] suggest its involvement in binding ions and small molecules, potentially influencing phage interactions with host bacteria (**Fig. 2k**). Given the role of cations in phage attachment to host cells[39], variations in this protein sequence may significantly impact virus-host interactions. Thus, our findings indicate that differences in viral infection proteins may contribute to varying bacteria-virus interactions across populations, reflecting unique viral adaptations.

### Predicted bacterial hosts of population-stratified viruses display distinct genetic components linked to various human health-related functions

Given the genomic stratification observed in predominant viruses and the role of mutations in bacterial infection-related proteins, we examined whether their bacterial hosts also exhibit genomic stratification. Using CRISPR spacer sequences from prokaryotic genomes, which serve as molecular records of past viral encounters[40, 41], we predicted bacterial hosts for the 44 VSCs showing population differences.

By comparing CRISPR spacer sequences from 2,791 virus genomes associated with the 44 VSCs to 478,588 global gut bacterial genomes[42], we identified 889,417 matches spanning 2,740 virus genomes and 17,897 bacterial genomes from 452 species. To enhance the confidence of these virus-bacteria pairs, we retained only those supported by at least 20 viral genomes within each VSC, resulting in 117 pairs involving 25 VSCs and 68 bacterial species (**Table S7**). Notably, the bacterial hosts in these pairs predominantly belonged to the phyla *Bacteroidota* and *Bacillota_A* (**Fig. 3a; Table S7**). Furthermore, comparisons of intra-group genomic dissimilarity among bacterial hosts corresponding to different viral sub-clades revealed significant differences in the genetic makeup of 23 bacterial host species based on their SNVs, as profiled by Snippy and analyzed using a Kimura 2-parameter distance matrix (FDR < 0.05; **Table S8**).

**Figure. 3.**
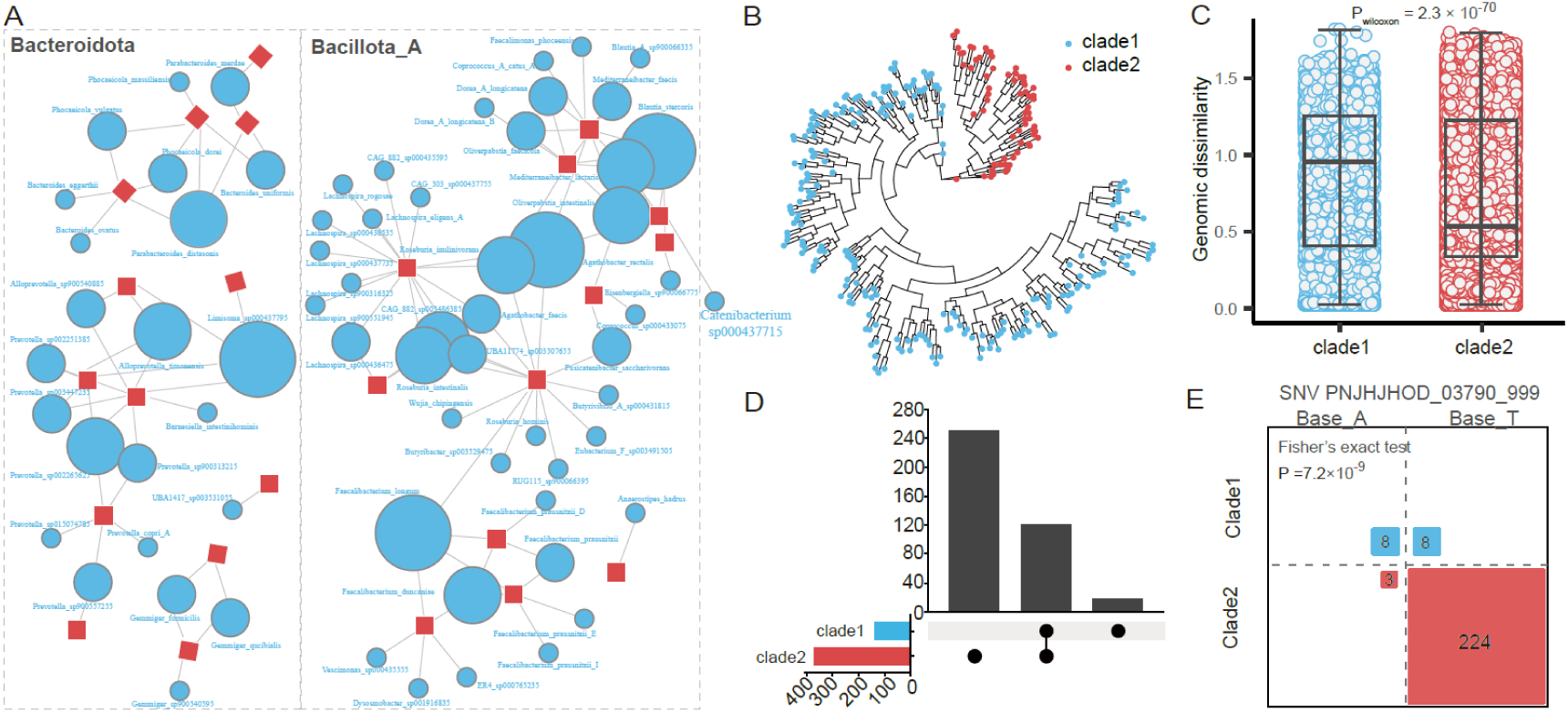
Predicted bacterial hosts of population-stratified viruses exhibit unique genetic components associated with diverse health-related functions including multidrug resistance. **a.** Predicted virus-bacteria pairs categorized by bacterial hosts at the phylum level. A total of 117 virus-bacteria pairs involves 25 VSCs and 68 bacterial species. Red indicates viruses, blue indicates bacterial hosts, and gray boxes represent bacterial hosts categorized by phyla, including *Bacillota_A* and *Bacteroidota*. **b.** Distinct sub-species clades of *FNMHLBE15_vae_111* genomes based on SNV profiles, with blue for clade 1 and red for clade 2. **c.** Genetic dissimilarity of the predicted bacterial host *Parabacteroides distasonis* within the two distinct sub-species clades of *FNMHLBE15_vae_111*, with blue and red indicating within-clade genomic dissimilarities. **d.** Summary of the predicted bacterial host *P. distasonis* specific to the sub-species clades of *FNMHLBE15_vae_111*. **e.** Differential allele types within the gene *PNJHJHOD_03790_999* (antibiotic ABC transporter ATP-binding and multidrug export protein) in *P. distasonis* between the sub-species clades of *FNMHLBE15_vae_111*.

To further characterize the genomic components driving the observed genetic stratification in bacterial hosts, we conducted an enrichment analysis to identify differential SNVs from bacterial genomes unique to various viral sub-species clades. This analysis revealed 257 differential SNVs across 6 bacterial species, corresponding to 242 CDSs (**Table S9**). Annotation of these CDSs using UniRef50[43] indicated a diverse range of encoded functionalities, including biosynthesis pathways for amino acids and short-chain fatty acids, as well as mechanisms related to drug resistance (**Table S10**).

For instance, 284 viral genomes from the *FNMHLBE15_vae_111* VSC formed two distinct sub-species clades (**Fig. 3b**). We predicted 390 metagenome-assembled genomes (MAGs) from *Parabacteroides distasonis* as their bacterial hosts based on CRISPR spacers, which exhibited significant genetic divergence (P_Wilcoxon_ = 2.3 × 10⁻⁷⁰) (**Fig. 3c; Table S8**). Notably, 18 MAGs were specific to one cluster, while 251 were specific to the other (**Fig. 3d**), and we identified 205 SNVs differentially enriched between the two groups. Among these, several SNVs were located in genes associated with drug resistance relevant to human health. In particular, we observed a significant differential enrichment of allele types at a locus within the *PNJHJHOD_03790_999* gene (Fisher’s exact test, P= 7.2 × 10⁻⁹) (**Table S9**), which encodes antibiotic ABC transporter ATP-binding[44] and multidrug export protein[45] (**Fig. 3e; Table S10**). These findings suggest the potential of leveraging population-specific viral strains to target bacterial hosts harboring antibiotic resistance genes, offering a promising strategy for combating antibiotic resistance through virus-mediated bacterial control.

### Within viral species genetic diversity is related to human phenotypic traits that independent of abundance variations

In addition to the genomic stratification of predominant viruses between populations, which suggests potential susceptibility to bacterial infections, we further examined whether within-species genomic variations are also associated with human host physiological traits. We filtered our cohort to identify VSCs for which viral genomes could be constructed from at least 10 MAGs, resulting in a total of 446 VSCs. Correlation analyses between the genetic dissimilarities of these VSCs and host phenotypic differences revealed 117 significant associations (FDR < 0.05) across 67 VSCs and 18 phenotypic traits (**Fig. 4a; Table S11**).

**Figure. 4.**
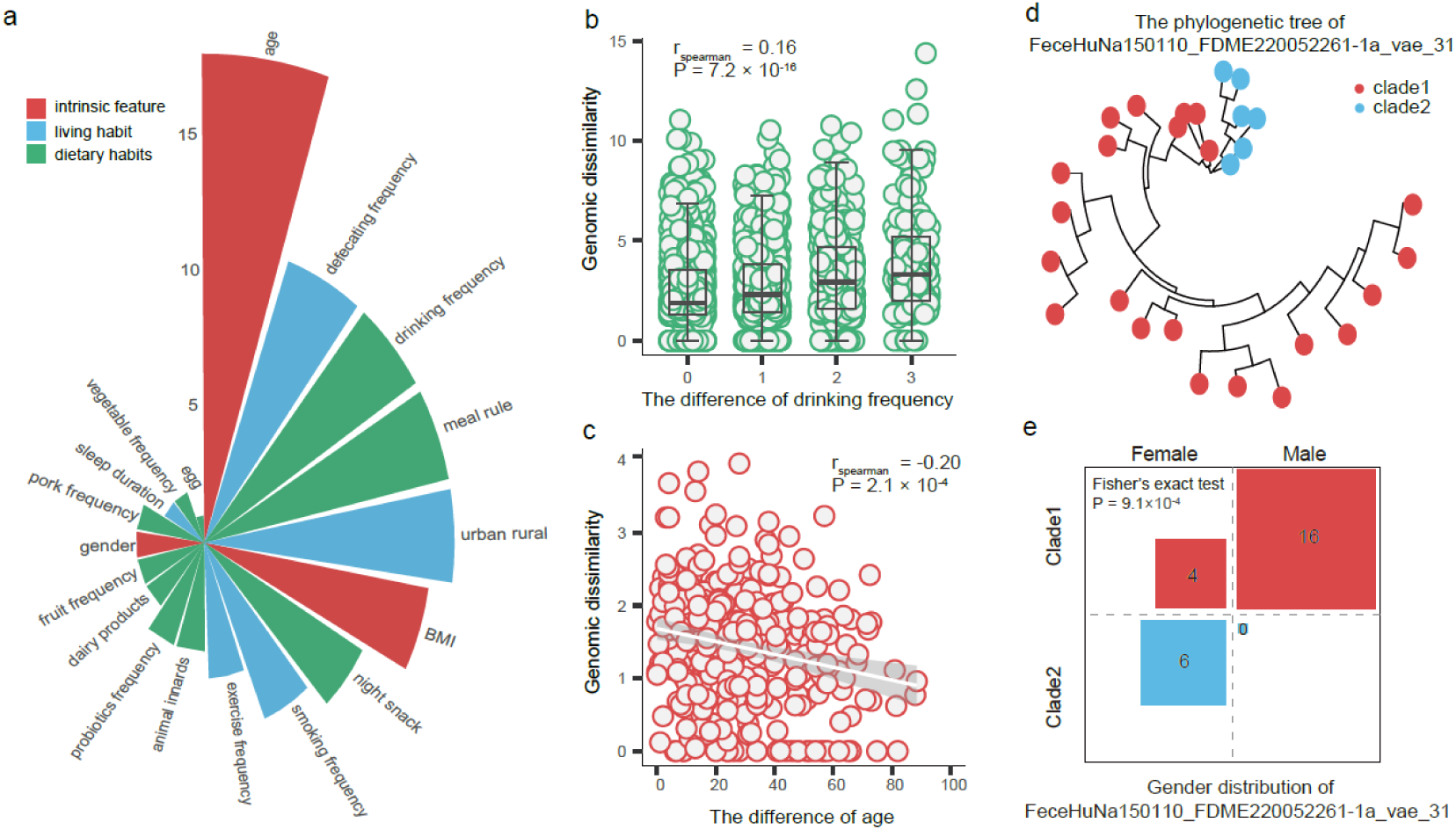
Genetic diversity within viral species is associated with host phenotypic traits. **a.** Summary of associations between VSC genomic dissimilarity and host phenotypic differences. Each bar represents a phenotype, sorted by the number of associations (FDR<0.05), with color coding: red for physiological traits, blue for lifestyle factors, and green for dietary habits. **b.** Genomic dissimilarity of *FSHCM57_vae_33* associated with drinking frequency. **c.** Genomic dissimilarity of *FGZ37M_FDME220052037-1a_vae_29* related to host age. **d.** Distinct sub-species clades of *FeceHuNa150110_FDME220052261-1a_vae_31* based on SNV profiles, with red indicating clade 1 and blue indicating clade 2. **e.** Gender distribution of hosts for *FeceHuNa150110_FDME220052261-1a_vae_31*, highlighting the differential distribution of viral genomes between male and female hosts.

Generally, within-species genetic dissimilarities were mainly linked to dietary habits and intrinsic factors like age and BMI (**Fig. 4a**). Notably, participants exhibiting greater phenotypic differences tended to show increased genetic variation in VSCs. For example, differences in drinking frequency were correlated with genetic variations in *FSHCM57_vae_33* (r_Spearman_= 0.16, P = 7.2×10⁻¹⁶) (**Fig. 4b; Table S11**). Additionally, many associations were observed with intrinsic factors such as age, BMI, and defecation frequency (**Fig. S3**). For instance, inter-individual age differences were associated with the genetic dissimilarities of *FGZ37M_FDME220052037-1a_vae_29* (r_Spearman_= –0.20, P = 2.1×10⁻⁴) (**Fig. 4c; Table S11**).

Given the substantial genetic diversity within viral species, we further investigated whether these viral genomes could form distinct sub-clades within the population. We hypothesized that main types of viral strains with similar genetic makeups might lead to unique functional attributes influencing host physiology. For 31 out of the 67 VSCs, distinct subspecies clades were observed in phylogenetic analyses (**Fig. S4**). By linking these viral subspecies clades to the aforementioned host phenotypic traits, we identified four significant associations between four VSCs and three host phenotypic traits (FDR < 0.05, **Table S12**). For instance, viral genomes from the *FeceHuNa150110_FDME220052261-1a_vae_31* VSC stratified into two clades specifically associated with differences in host sex (P = 9.1×10⁻⁴, **Fig. 4d, e**).

To determine whether the observed genetic associations between VSCs and host phenotypic traits were influenced by abundance differences, we filtered for VSCs present in at least 20% of participants and comprising a minimum of 10 viral genomes, yielding 408 VSCs. We profiled their abundance in fecal metagenomic samples using transcripts per kilobase of exon model per million mapped reads (TPM) calculated by CoverM. Among these VSCs, 87.0% (355 out of 408) showed no significant association between genomic dissimilarity and abundance differences (FDR > 0.05, **Fig. S5; Table S13**), suggesting that genetic variability among VSCs provides additional insights independent of their abundance. These findings underscore the critical role of viral strains in influencing host phenotypic traits.

### Abundance variations of newly characterized VSCs widely associated with human phenotypic traits

As demonstrated, within-species viral genomic variations largely operate independently of abundance differences, with only modest associations observed. To provide further biological relevance for the newly characterized VSCs lacking taxonomic information, we linked their abundances to human phenotypic traits. We profiled the viral abundance of 38,436 unique VSCs in fecal metagenomic samples, detecting an average of 27,310 VSCs per sample and 35,549 VSCs in over 20% of samples.

Using MaAsLin2[46], a tool for analyzing multivariable associations between host phenotypes and microbiomes, we identified 19,108 significant associations between 13,215 VSCs and 22 phenotypes (FDR < 0.05), primarily enriched for geographic, individual intrinsic, and dietary factors (**Fig. 5a; Table S14**). Notably, residential location (urban vs. rural), defecation frequency, and age were the most strongly associated with viral abundance.

**Figure. 5.**
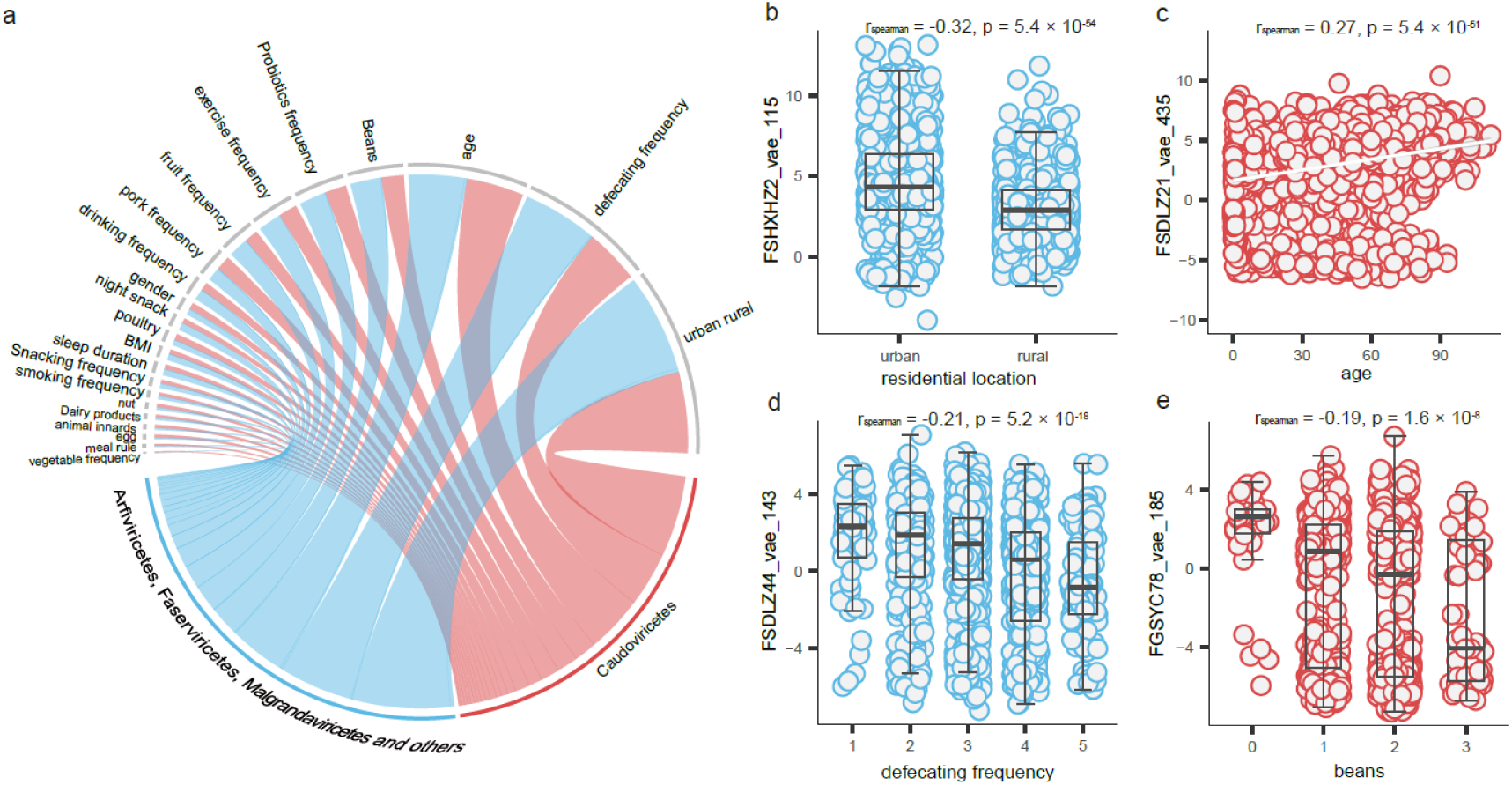
Abundance variations of newly characterized VSCs widely associated with human phenotypes. **a.** Summary of associations between VSC abundances and various phenotypic traits. Human phenotypes are represented in gray, while gut viruses from different viral classes are color-coded. Each line indicates a significant association between a viral factor and a phenotype, with line colors corresponding to the associated viral class. **b.** The relative abundance of *FSHXHZ2_vae_115* is significantly higher in urban populations compared to rural areas. **c.** The relative abundance of *FSDLZ21_vae_435* shows a positive correlation with host age, indicating that older individuals tend to harbor higher levels of this viral species. **d.** The relative abundance of *FSDLZ44_vae_143* is negatively associated with defecation frequency, suggesting that increased levels of this viral species may be linked to lower frequency of bowel movements. **e.** The relative abundance of *FGSYC78_vae_185* is negatively associated with the frequency of bean consumption, indicating that individuals who consume beans more frequently tend to have lower levels of this viral species.

For instance, participants in urban areas exhibited a higher abundance of *FSHXHZ2_vae_115* (r_spearman_ = –0.32; P = 4.4 × 10⁻⁵⁴, **Fig. 5b**), while the abundance of *FSDLZ21_vae_435* increased with age (r_spearman_ = 0.27; P = 5.4 × 10⁻⁵¹, **Fig. 5c**). Additionally, variations in defecation frequency correlated with viral abundance, as observed with *FSDLZ44_vae_143*, which showed a negative association with increased defecation frequency (r_spearman_ = –0.21; P = 5.2 × 10⁻¹⁸, **Fig. 5d**). Regarding dietary habits, the frequency of bean consumption was negatively associated with *FGSYC78_vae_185*, a member of *Caudoviricetes* (r_spearman_ = –0.19; P = 1.6 × 10⁻⁸, **Fig. 5e**). These findings enhance our understanding of the potential roles of taxonomically unclassified gut viruses in human phenotypic traits, elucidating their potential roles in human health and disease.

### Inter-individual gut bacteriome abundance differences is specifically shaped by virome

The human gut virome is primarily dominated by bacteriophages, which regulate bacteria by either lysing them or modifying their physiological functions[47]. Among the 20,220 out of 38,436 VSCs with taxonomy information available in our CGVR, 90.3% (18,253) were identified as bacteriophages (**Table S15**). To assess the extent to which inter-individual variations in the gut bacteriome are shaped by the virome, we profiled the abundance of 5,310 bacterial species using MetaPhlAn4. Our analysis revealed a positive correlation between the number of identified viruses and the number of detected bacterial species per sample (**Fig. 6a**).

**Figure. 6.**
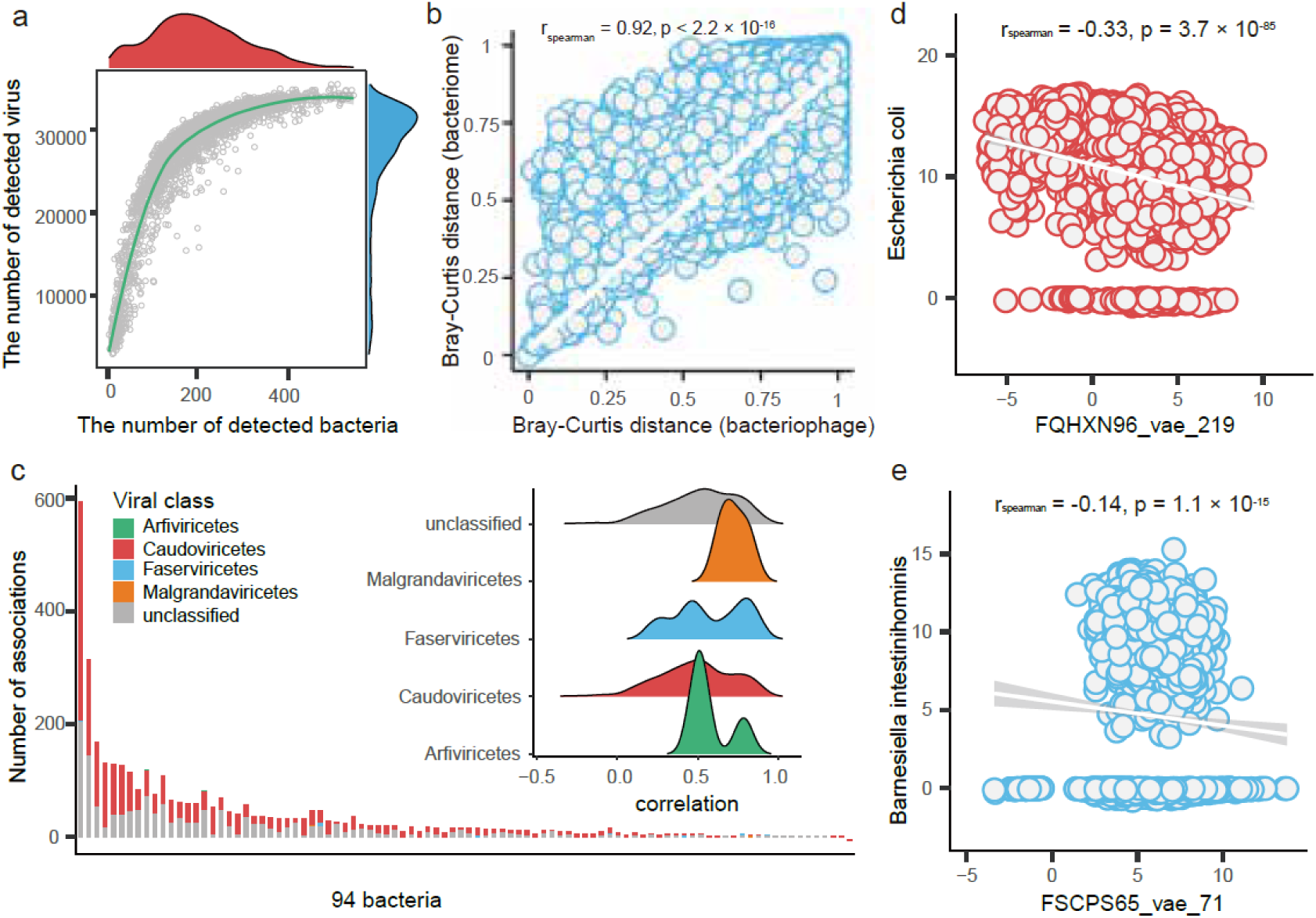
The gut bacteriome composition variations are specific to virome. **a.** Relationship between the number of identified viruses and the number of detected bacterial species per sample. Red density represents the number of detected bacteria, while blue density represents the number of detected viruses. Each dot indicates an individual. **b.** Correlation of overall virome and bacteriome composition based on Bray-Curtis distance calculated using abundance data. Each dot represents the Bray-Curtis distance between pairs of individuals. Regression lines are shown in white. **c.** Summary of viral abundance associations with bacterial abundance confirmed by CRISPR-spacer imprints. The bar plot summarizes the number of VSCs with significant associations per bacterial species, while the histogram shows the distribution of correlation coefficients based on viral class. **d.** Negative association between *FQHXN96_vae_219* and *Escherichia coli*abundances. **e.** Negative association between *FSCPS65_vae_71* and *Barnesiella intestinihominis* abundances.

By further calculating the Bray-Curtis distance matrix based on the abundance of 18,253 VSCs and 5,310 bacteria, we found a strong correlation between the two matrices (r_spearman_ = 0.92, P < 2.2 × 10⁻¹⁶, **Fig. 6b**). Notably, a total of 84.3% of the bacterial compositional variance can be attributed to the gut virome composition estimated by linear regression, indicating that the human gut bacteriome composition is largely shaped by the virome.

In addition to analyzing the overall gut virome and bacteriome composition, we identified 41,844 significant associations involving the abundance of 6,131 VSCs and 302 bacterial species using the MaAsLin2 (FDR < 0.05, **Table S16**). These associations were primarily enriched in the phyla *Pseudomonadota*, *Bacillota*, *Actinobacteria*, and *Bacteroidota*. To assess the concordance with CRISPR-spacer-based analyses, we focused on significant associations that corresponded with CRISPR-spacer imprints, resulting in a total of 3,497 associations (**Table S17**, **Fig. 6c**). Among these, 2.1% were negative (**Fig. 6c**), indicating that lower bacterial abundance was linked to higher viral abundance, suggesting potential viral regulation of bacterial composition.

At the individual species level, the most significant viral associations were observed with *Escherichia coli* (**Fig. S6**). For instance, we found that a lower abundance of *E. coli* was associated with a higher level of *FQHXN96_vae_219* (r_spearman_ = –0.33; P = 3.7 × 10⁻⁸⁵, **Fig. 6d**). The emergence of antibiotic-resistant *E. coli* has become a global concern due to the widespread use of antibiotics[48–50]. Our results suggest the potential for targeted lysis of *E. coli* cells by specific viruses, thereby addressing antibiotic resistance and infections related to *E. coli*[51].

Furthermore, our data revealed associations between the abundance of probiotics and viruses. Probiotics like *Barnesiella intestinihominis* have been shown to enhance immune responses, improving the effectiveness of chemotherapy and immunotherapy[52]. In our findings, higher abundances of *B. intestinihominis* were associated with lower abundances of *FSCPS65_vae_71* (r_spearman_ = –0.14; P = 1.1 × 10⁻¹⁵, **Fig. 6e**). These results further underscore the potential of targeting gut viruses to regulate bacterial populations related to human health and disease.

### The gut viral abundance associations to human phenotypic traits are mediated by bacteriome

Since the abundance of the gut virome can be linked to both the bacteriome and human phenotypic traits, and given that gut viruses like bacteriophages are unlikely to interact directly with human hosts, the viral impact on human phenotypic traits may be mediated by the bacteriome. To evaluate this, we conducted mediation analysis focusing on 966 viruses associated with both the bacteriome and human phenotypic traits (**Fig. 7a**). We identified 107 significant mediation linkages involving 88 viral species (FDR < 0.05, **Table S18**), the main associations are related to individuals’ basic characteristics and lifestyles (**Fig. 7b**).

**Figure. 7.**
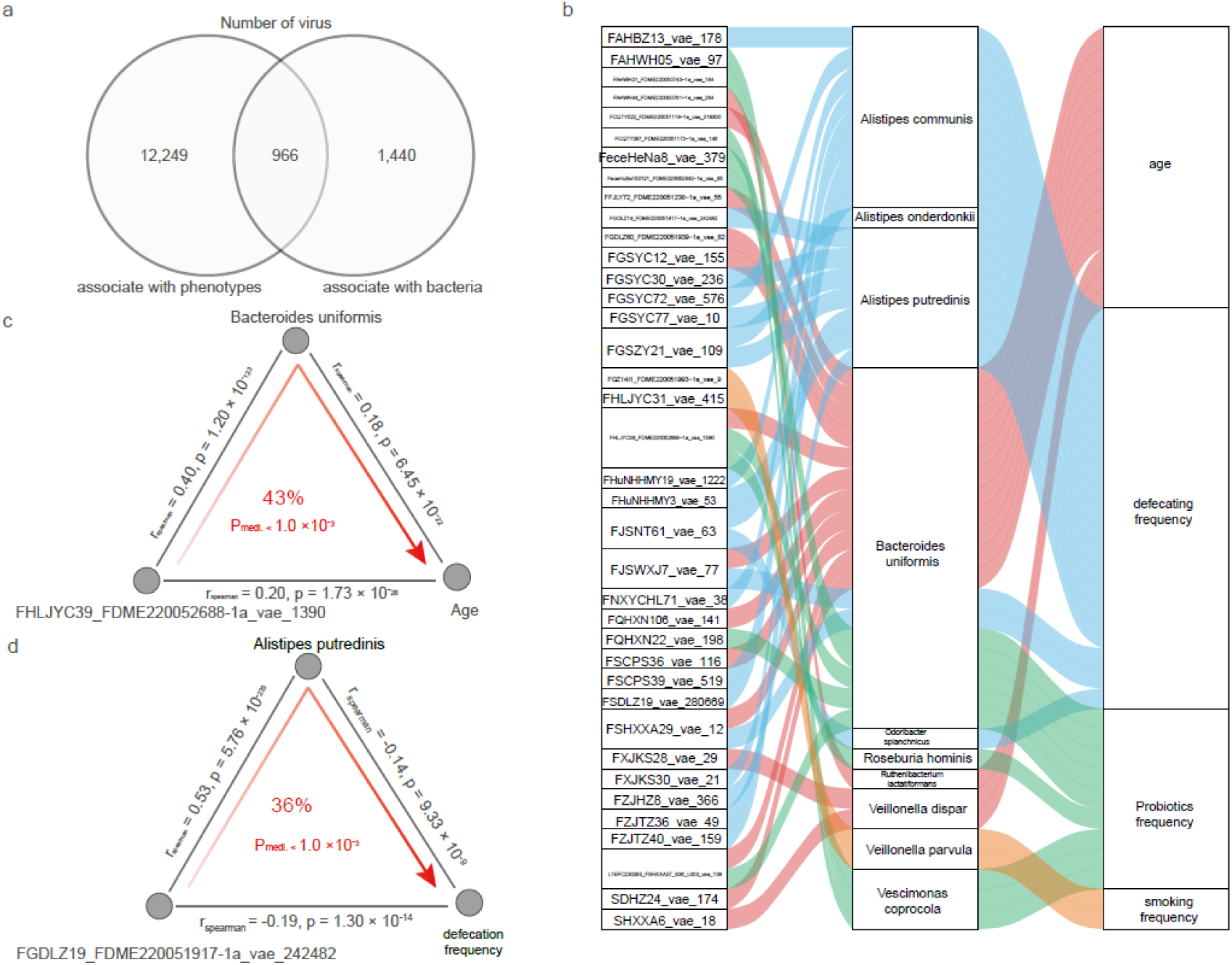
Mediation linkages among the gut virome, bacteriome, and human phenotypes. **a.** Venn diagram showing the number of viruses associated with phenotypes and bacteria, respectively. **b.** Parallel coordinates chart displaying 107 significant mediation effects of gut microbial species. The left panel shows the viral species (VSCs), the middle panel presents the microbial species, and the right panel illustrates the human phenotypes. Curved lines across the panels indicate mediation effects, with colors representing different phenotypes. **c.** Microbial species *Bacteroides uniformis* mediates the effect of viral species *FHLJYC39_FDME220052688-1a_vae_1390* on age. **d.** Microbial species *Alistipes putredinis* mediates the effect of viral species *FGDLZ19_FDME220051917-1a_vae_242482* on defecation frequency.

Most of these linkages were related to the viral impact on host phenotypes via *Gemmiger formicilis* (19 linkages) and *Bacteroides uniformis* (18 linkages), highlighting the profound influence of viruses on the complex host-microbe ecosystem in the intestine. For instance, the abundance of *B. uniformis* is positively correlated with age, and the virus *FHLJYC39_FDME220052688-1a_vae_1390* demonstrates a symbiotic relationship with *B. uniformis*, mediating its abundance (43%, P_mediation_ < 0.001, **Fig. 7c**). Additionally, the abundance of virus *FGDLZ19_FDME220051917-1a_vae_242482* with a decrease in defecation frequency, a phenotypic trait that relevant for gastrointestinal issues, which correlates with a reduction in *Alistipes putredinis* (36%, P_mediation_ < 0.001, **Fig. 7d**). Taken together, these findings provide potential mechanistic underpinnings for virome-microbiome interactions in human phenotypes.

## DISCUSSION

In this study, we present the Chinese Gut Viral Reference (CGVR) set, comprising 120,568 viral genomes from 3,234 fecal samples collected nationwide. Through sequence clustering, we identified 72,751 viral species-level clusters (VSCs), with nearly 90% absent from existing databases[6, 9–19]. We found population-specific genomic differences in common gut viral species between Chinese and non-Chinese populations, which are linked to bacterial host infections. Additionally, the predicted bacterial hosts of these population-stratified viruses exhibit distinct genetic components associated with various health-related functions, such as multidrug resistance. Our analysis revealed that within-species viral genomic variations correlate with host physiological traits at the single-nucleotide level, independent of abundance variations. Moreover, we observed that inter-individual differences in the gut bacteriome are specifically shaped by the virome, identifying 19,108 viral abundance associations with host phenotypes, enriched in geographic, intrinsic, and dietary factors. Finally, mediation analysis suggests that the gut virome may influence human phenotypic traits through interactions with the bacteriome.

The human gut virome is a critical part of the microbiome and characterizing viral genomic diversity and functional variations across global populations is crucial in addressing regional health disparities. However, large-scale construction of viral genomes has primarily focused on samples from western populations[6, 9–19], hindering a full understanding of global gut viral genomic diversity and functional variations at strain-level resolution. To address this limitation and expand the global reference of viral genomes, we compared our Chinese dataset with the UHGV dataset, adding 38,436 novel species-level clusters to existing reference genomes. This significantly broadens the representation of viral genomes, particularly for non-Western populations. Our analysis of viral species-level clusters (VSCs) revealed significant population-specific stratification. Previous studies have highlighted the geographic variability of viral species[6, 20], emphasizing the vast diversity of the human gut virome and the necessity for strain-resolved genome databases for functional interpretation.

The gut virome is a crucial component of the microbiome, and understanding virus-bacteria interactions is vital[15, 53]. While prior studies linked the virome to bacterial hosts[6, 10, 30], they often overlooked population-specific features of the gut virome. This is essential for comprehending the mechanisms of virus-specific infections. Using the CRISPR spacer method, we identified that bacterial hosts of population-stratified viruses possess distinct genetic elements linked to health-related functions. Furthermore, our findings show that within-species viral genomic diversity correlates with host phenotypic traits independent of abundance variations.

In addition to genetic diversity, viral abundance plays a key role in human health and disease. Through associations between newly characterized VSCs and bacteria, we identified tens of thousands of potential virus-phenotype and virus-bacteria associations. Mediation analysis revealed over a hundred significant associations between gut viral abundance and human phenotypic traits mediated by the bacteriome. These findings highlight the potential of targeting gut viruses to modulate bacterial populations relevant to human health.

Notably, while previous studies demonstrated the role of viruses in regulating bacterial populations and metabolites through experimental evidence[54], our results, based on a large-scale dataset, offer lots of but observational insights into the virome’s role in human health, which may provide valuable direction for future research.

We acknowledge several limitations in our study. First, while we collected over 3,000 samples from 30 provinces, the sample size remains relatively small compared to China’s population. Second, current virus detection tools may have limitations, leading to inaccuracies in identifying viral sequences[55, 56]. Third, advanced virome analysis techniques, such as long-read metagenomic sequencing or virus-like particle sequencing, have successfully uncovered hidden diversity in human gut viruses[7, 53]. Employing these methods could provide a more comprehensive view of the virome in the Chinese population.

Taken together, our study expands the existing collection of gut viral genomes and offers functional insights into genetic diversity and population specificity of the human gut virome at the single-nucleotide level. This work enhances our understanding of host-microbe interactions and lays the groundwork for developing microbiome-based personalized therapies.

## MATERIALS AND METHODS

### Human subjects

The study was approved by the Ethics Review Board of Jiangnan University (reference number JNU20220901IRB01) and included 3,234 participants from 30 provinces across mainland China. Informed consent was obtained from all participants prior to sample collection. Comprehensive phenotypic data were gathered, encompassing anthropometric traits (e.g., age, sex and body mass index), geographical factors, residential location (urban or rural), dietary habits, lifestyles, as well as medical history and medication use. A detailed summary of these phenotypic traits has been previously reported[26].

### Sample collection and metagenomic sequencing

Fecal samples were collected and placed on dry ice within 15 minutes of production. The samples were then transported to the laboratory, where aliquots were prepared using a 25% pre-reduced anaerobic glycerol solution and 1% cysteine. The aliquots were stored at –80°C until further processing. DNA extraction from the fecal samples was carried out using the QIAamp Fast DNA Stool Mini Kit (Qiagen, Cat. 51604). After DNA extraction, library preparation and whole-genome shotgun sequencing were performed using the MGI DNBSEQ-T7 platform for 3,082 samples and the Illumina NovaSeq 6000 platform for 152 samples[26].

### Sequence preprocessing and assembly

All metagenomic data underwent quality control and decontamination of human genomic DNA (NCBI37) using KneadData (v0.12.0) with default parameters. The following key steps were employed to ensure high-quality datasets and assembly of reads into contigs: (i) trimming of adapters and low-quality bases, (ii) identification and masking of human reads, and (iii) assembly of clean reads. The raw reads were trimmed to remove adapters and low-quality bases at both ends using Trimmomatic[57] (v.0.39). Reads shorter than 50 bp after trimming were discarded. Subsequently, the quality-filtered reads were aligned to the human reference genome using Bowtie2[58] (v.2.4.2). Reads for which both paired ends failed to align were retained. After completing data quality control, MegaHIT[59] (v.1.2.9) was used to assemble the quality-filtered reads, generating 6.9 × 10⁷ contigs.

### Viral genomic binning and quality assessment

Viral genomic binning was performed using VAMB[60] (v.4.1.3), a deep-learning-based tool that clusters metagenomic contigs into putative biological entities based on assembly sequences. During this process, contigs shorter than 2 kb were excluded to ensure data quality. After binning, all VAMB-generated bins were further refined using the viral binning method PHAMB[61] (v.1.0.1), resulting in the identification of approximately 5.5 × 10⁶ putative viral bins. To assess the completeness of the metagenome-assembled viral genomes, we applied CheckV[27] (v.1.0.1, database v1.5) with the “end_to_end” mode and default settings. This step ensured that only high-quality viral genomes were included in further analyses, contributing to a robust dataset for viral characterization.

### Generation of viral species-level sequence clusters (VSCs)

To generate species-level viral sequence clusters (VSCs), we first built a nucleotide BLAST database of all 64,377 viral sequences using makeblastdb (option: “-dbtype nucl”) from BLAST[62] (v.2.14.1). Pairwise comparisons of the viral sequences were generated using an all-vs-all local alignment strategy, executed via blastn with the option “-max_target_seqs 10000”. Next, two custom scripts, anicalc.py and aniclust.py, from the CheckV repository’s genome clustering tools, were employed to compute average nucleotide identity (ANI) and alignment fraction (AF) for clustering sequences at the species level. Using the anicalc.py script, pairwise ANI values were calculated by merging local alignments between sequence pairs. Finally, species-level clustering was performed using aniclust.py in a UCLUST-like manner with the recommended parameters (“-min_ani 95, –min_tcov 85, –min_qcov 0”) from the MIUVIG[28] protocol, effectively grouping viral genomes into species-level clusters.

### Taxonomic annotation of viral genomes

Each viral genomes were taxonomically classified using uhgv-tools (v0.0.1) with ‘classify’ mode. The uhgv-tools utilize aligning genome and protein sequences against an integrated viral database (uhgv-db-v0.4) to obtain average DNA identity, alignment fraction and amino acid similarity scores, which are used to propose viral ICTV^56^ taxon lineage.

### Functional annotation of genes in viral genomes

The open reading frames (ORFs) of the viral genomes were predicted using Prodigal[31] (v2.6.3) with the option “-p meta”. Functional annotation of the genes was performed using eggNOG-mapper[32] (v2.1.12), utilizing the EggNOG database[33] (v5.0.2) with default settings. The resulting functional annotations included categories from COG, KEGG[63], and Pfam[64], which were derived from the eggNOG-mapper output. Eleven classes of phage parts were categorized based on their functions, including LYS (lysis), INT (integration), REP (replication), REG (regulation), PAC (packaging), ASB (assembly), INF (infection), EVA (immune evasion), HYP (hypothetical protein), UNS (unsorted), and tRNA according to a previous study[65].

### Comparison of viral genomes with the UHGV

Viral sequences from the human gut virome catalog database (UHGV, https://github.com/snayfach/UHGV) were obtained for comparison. The UHGV database integrates gut virome collections from 12 recent studies[6, 9–19], including MGV, GPD, GVD, GFD, CHVD, Hadza, Benler, mMGE, DEVoC, IMG/VR, HuVirDB and COPSAC, totaling 2,242,702 putative viral genomes. To ensure data quality, the completeness of all viral sequences was assessed using CheckV. From this assessment, 208,604 high-quality viral sequences were selected for comparative analysis. A nucleotide BLAST database of all quality-controlled viral sequences was then constructed, and pairwise comparisons were performed using all-vs-all local alignments with blastn (v2.14.1).

### Single nucleotide variant (SNV) calling

For each cluster of selected viruses, all protein-coding genes of VSCs were predicted using Prodigal (v2.6.3) with the parameters –p meta –f gff to generate GFF files. Subsequently, we utilized Roary^59^ (v3.12.0) with options –s –i 95 to obtain a pan-genome reference, which serves as a comprehensive and reliable reference for each specific virus species. To detect SNVs within each VSC, we used the pan_genome_reference.fa file generated by Roary as the reference genome. The detection process was performed using Snippy (v4.6.0), a fast and accurate tool specifically designed to identify single nucleotide polymorphisms (SNPs) and insertion-deletion events (indels) in draft or complete genomes through a mapping-based approach. A coverage threshold of 10 reads at a site was established to consider an SNV as valid. The SNVs of bacterial hosts were characterized using the same methodology.

### Genomic dissimilarity

For each viral and bacterial genome, we focused on the single nucleotide polymorphism (SNP) variants extracted from the Snippy output, temporarily excluding insertions, deletions, and complex variants. We combined the .vcf files of all genomes associated with the same species into a matrix format, where each row represented a variant site and each column corresponded to the nucleotide type of a genome at that site. Each column was treated as a distinct sequence, which we aligned to generate a phylogenetic distance matrix. Pairwise nucleotide substitution rates between samples were calculated using the Kimura 2-parameter method from the EMBOSS[66] package. For any NA values in the distance matrix, indicating distances that were too large to compute, we replaced these entries with the maximum distance value plus one.

### Phylogenetic trees

To identify distinct sub-species clades or strains, we employed hierarchical clustering using the complete linkage method. This approach was applied to a genomic distance matrix derived from SNP haplotype data, enabling the classification of viral and bacterial strains based on their genetic similarity. Hierarchical clustering of genomes was performed using *hclust* function. Phylogenetic trees were then visualized using *ggtree*^60^ package (v3.6.2).

### Protein structure and function prediction

To investigate the three-dimensional structures and potential functions of the virus proteins of interest, we employed advanced computational tools, including AlphaFold3[37] and DeepFRI[38]. After predicting protein structures using AlphaFold3, we utilized DeepFRI to infer potential functions. DeepFRI employs a pre-trained graph-based neural network model to analyze protein sequences and structures, providing a list of predicted functions along with confidence scores for each protein. Protein structures were visualized using PyMOL.

### CRISPR spacer-based prediction of viral bacterial hosts

To predict bacterial hosts for VSCs, we utilized the Global Microbiome Reference (GMR) catalog[42] to identify CRISPR spacers. This was achieved by integrating the outputs from two CRISPR detection tools, CRT[67] and PILER-CR[68], both executed with default settings. The predicted CRISPR spacers were then aligned against the representative VSCs from the CGVR catalog using blastn (v2.14.1). The alignment was optimized for short sequences with the following options: –max_target_seqs 10000, – task blastn-short, –evalue 1, –gapopen 10, –gapextend 2, –penalty –1, –word_size 10, and –perc_identity 100. This approach allowed for accurate matching of CRISPR spacers to viral genomes, facilitating bacterial host prediction.

### Functional annotation of genes in bacterial genomes

To investigate the functions of SNVs derived from metagenome-assembled genomes (MAGs) that contribute to the observed genetic stratification in bacterial hosts, we performed functional annotation of bacterial proteins using the UniRef50[43] database. This process involved conducting a BLAST alignment of the gene sequences against the UniRef50 database using Diamond with the parameters ‘--evalue 1e-5 –-id 50’.

### Relative abundance of VSCs

The TPM (transcripts per kilobase of exon model per million mapped reads) values were used to represent relative abundances of VSCs in the fecal metagenomic samples estimated by CoverM (v0.6.1). The representatives of VSCs (n = 42,803) were utilized to construct the mapping reference genome through bowtie2-build (version 2.4.2). Subsequently, the quality-controlled reads from each individual sample were mapped using bowtie2. Finally, the TPM values were computed with the “--methods tpm” option, leveraging custom scripts.

### Relative abundance of bacterial species

Bacterial species abundances were generated using MetaPhlAn4[69], which leverages approximately 5.1 million unique clade-specific marker genes identified from around 1 million microbial genomes. This extensive database encompasses 26,970 species-level genome bins, facilitating unambiguous taxonomic assignments, accurate estimation of organismal relative abundance, and species-level resolution for bacteria. Only microbial species present in more than 20% of the samples were included for further analysis, resulting in a list of 303 species.

### Quantification and statistical analysis Differential comparison

Viral genomic dissimilarities within and between Chinese and non-Chinese populations were assessed using the Kruskal-Wallis test, while genomic dissimilarities between the two groups were evaluated using the Wilcoxon test. P-values were adjusted using the Benjamini-Hochberg (BH) method[70].

### Enrichment of allele type

Differential enrichment of specific allele types in viral genomes between populations and sub-species clades was assessed using Fisher’s exact tests. P-values were adjusted using the BH method.

### Associations of within-species viral genomic dissimilarities with phenotypic differences

Spearman correlation was applied to assess whether genomic variations within viral species are associated with human host physiological traits. For inter-individual gut viral genomic dissimilarities, pairwise nucleotide substitution rates between samples were calculated using the Kimura 2-parameter method. The Manhattan dissimilarity metric was applied to measure phenotypic differences. P-values were adjusted using the BH method.

### Associations between vial genomic and abundance differences

Spearman correlation was applied to evaluate whether genomic variations within viral species are associated with abundance differences. For inter-individual gut viral genomic dissimilarities, pairwise nucleotide substitution rates were calculated using the Kimura 2-parameter method. The Manhattan dissimilarity metric was used to assess abundance differences, and P-values were adjusted using the BH method.

### Viral abundance associations with human phenotypes and bacterial abundance

The relative abundances of VSCs and bacterial species were centered log-ratio transformed and then subjected to Spearman correlation. No transformation was applied to the phenotypic data. For associations identified at FDR < 0.05, we conducted MaAsLin2 analysis to estimate relationships between viral features, human phenotypes, and bacterial species abundance. Human phenotypes and bacterial species significantly associated with VSCs were treated as fixed effects in MaAsLin2, and results with FDR controlled at 0.05 were reported.

### Association between virome and gut bacteriome overall compositions

We applied the *vegdist* function from the vegan (version 2.6.4) R package to calculate the Bray-Curtis dissimilarity matrix based on the relative abundances of the virome and bacteriome at the species level. We then assessed the correlation between the two distance matrices using the Spearman correlation test. To estimate the variance in bacterial overall composition explained by the gut virome, linear regression was applied.

### Mediation linkage inference

For VSCs associated with both human phenotypic traits and bacterial species, we first checked whether host phenotypic traits were also associated with bacterial species using both Spearman correlation and MaAsLin2 (FDR < 0.05). Next, mediation analysis was carried out using the mediate function from the R package mediation (version 4.5.0) to infer the mediation effect of bacterial species on viral associations with human phenotypic traits. P-values were adjusted using the BH method.

### Data availability

The metagenomic sequencing data used for the analysis presented in this study are available from the China National Gene Bank Data Base (CNGBdb) under accession id CNP0004122. The genomic assemblies of gut virus are deposited in National Genomics Data Center (NGDC) under accession id OMIX007646. Access to data should be subjected to the policies from the Human Genetic Resource Administration, Ministry of Science and Technology of the People’s Republic of China.

### Code availability

Scripts used for the analyses in this manuscript are available via: https://github.com/MicrobiomeCardioMetaLab/CGVR_project.

## ACKNOWLEDGEMENTS

We thank the participants and management staffs for their collaboration and support.

## AUTHOR CONTRIBUTIONS

L.C., X.K. and Q.Z. contributed to conceptualization and funding. The CGVR Consortium Initiative staffs contributed to data and sample collection. X.W., Q.D., L.C., P.H., S.Y. contributed to data analysis. Q.D., X.W., L.C. drafted the manuscript. X.W., Q.D., P.H., S.Y., M.G., C.Z., C.Z., Y.D., Z.H., B.M., Y.J., Y.Z., T.W., H.Z., J.S., Y.S., Y.W., L.T., S.H., Y.D., W.S., W.C., Q.Z., X.K. and L.C. contributed to discussion of the content. All authors read, revised and approved the final manuscript.

## Fig. S1 – S6

**Fig. S1.**
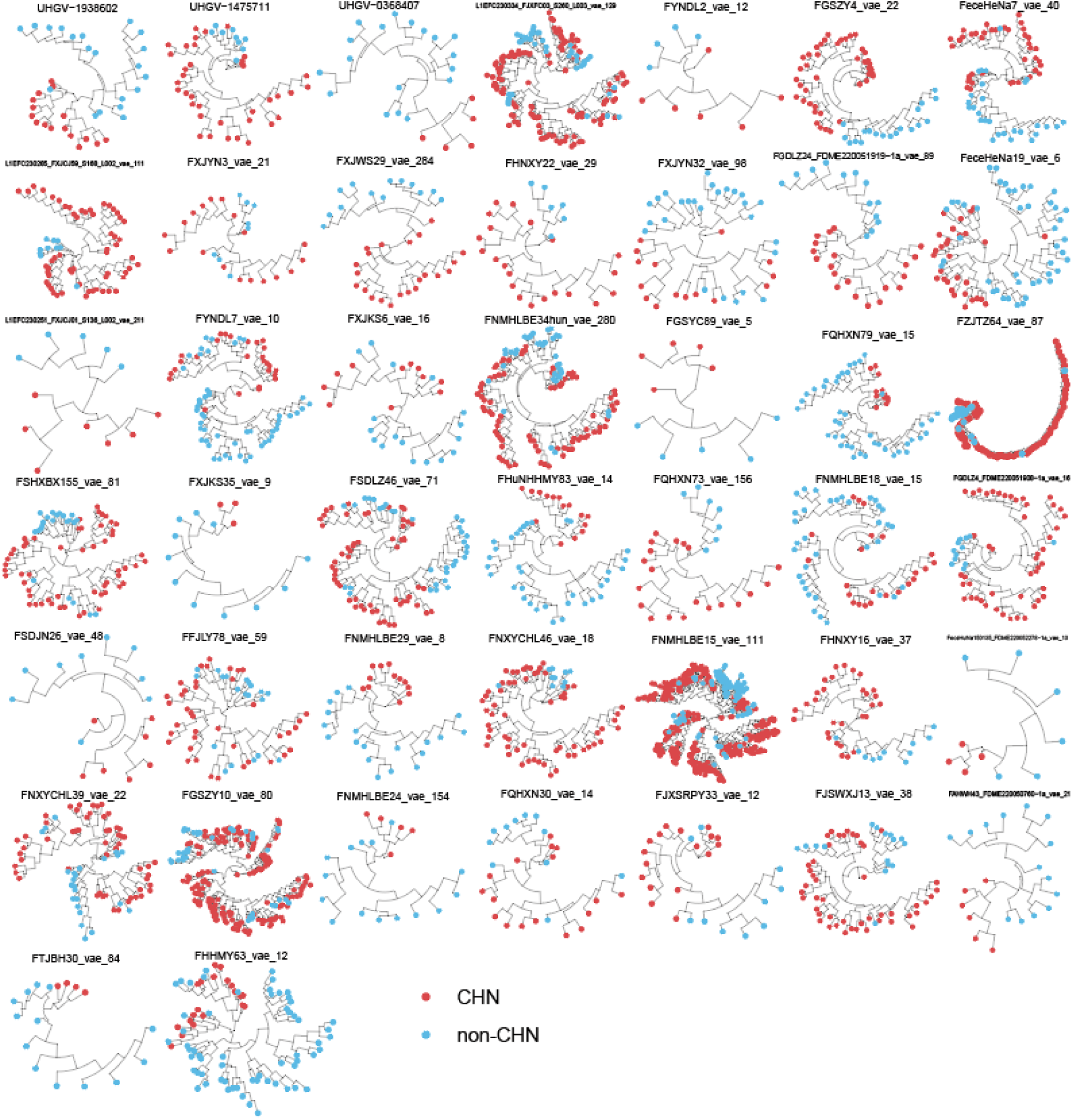
Phylogenetic tree of viral genomes from 44 VSCs based on SNP haplotype profile. Red represents viral genomes from Chinese population and blue represents the non-Chinese population.

**Fig. S2.**
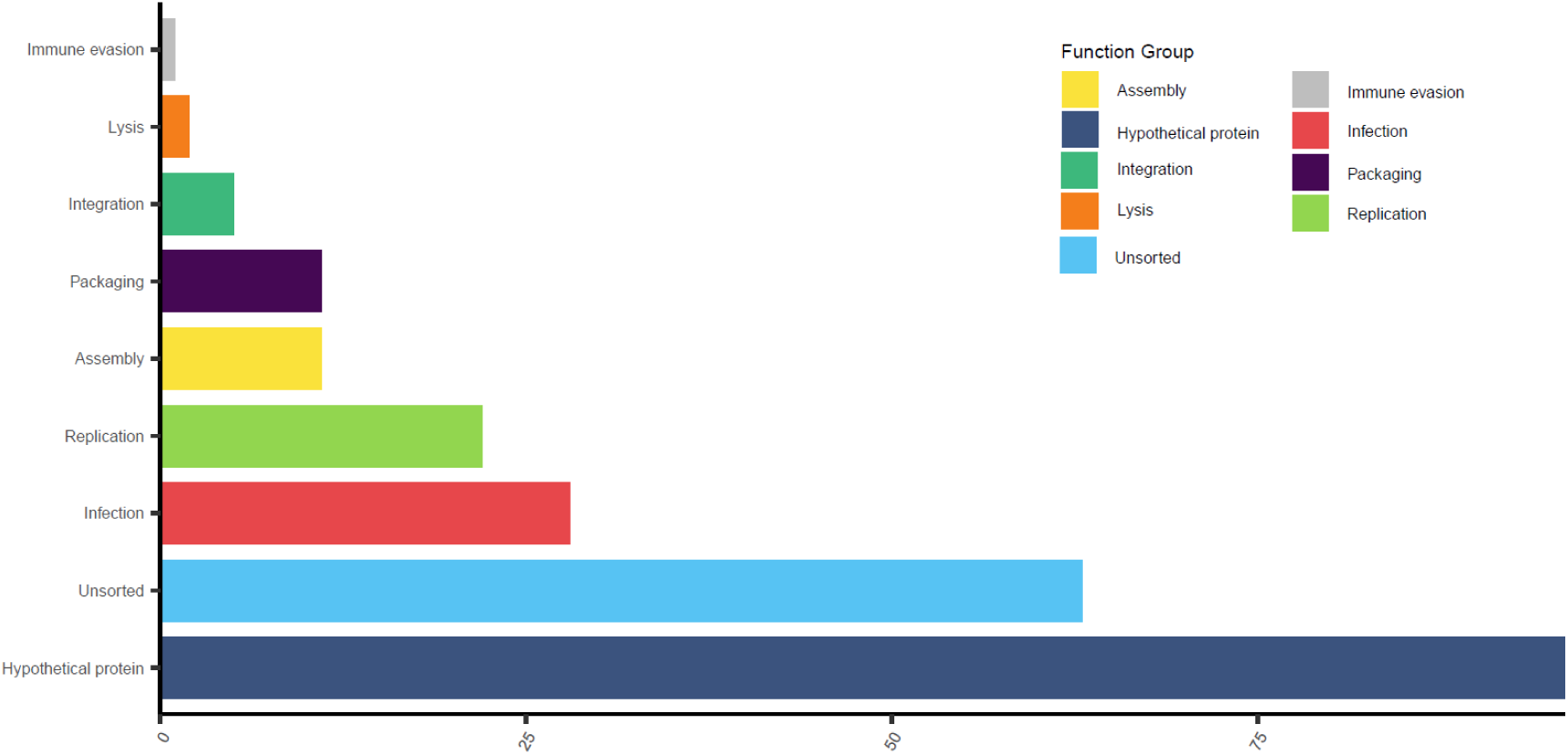
Functional annotation of viral genes with differential SNVs between populations. We conducted functional annotation of the 239 SNVs that exhibit significant population differences and classified them into different functional groups based on a previous study[65].

**Fig. S3.**
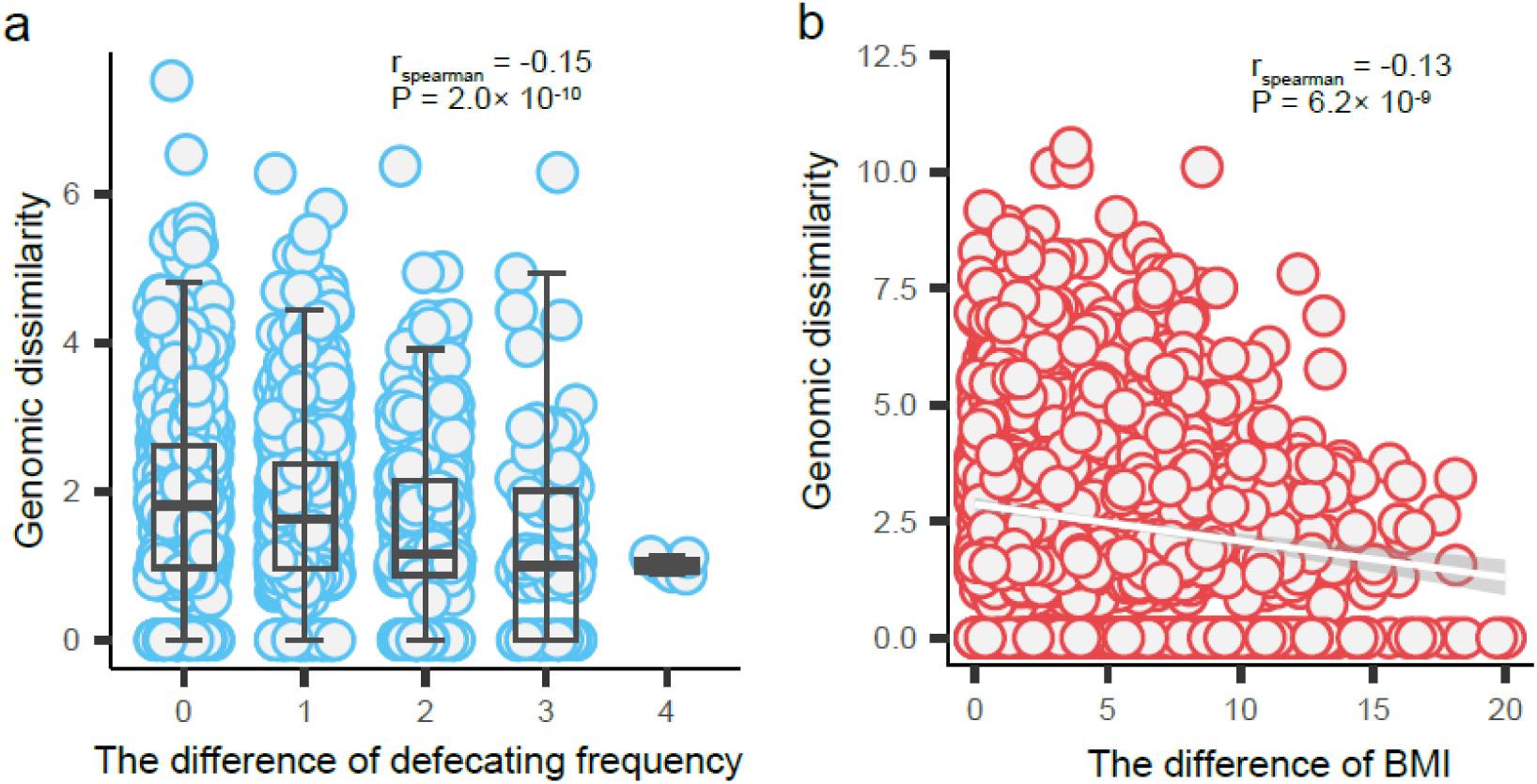
Associations between viral species genomic and host phenotypic differences. A. The genetic distance of L1EFC230390_FSHXXA28_S33_L003_vae_13 decrease with increasing variation in defecating frequency. B. The genetic distance of FSHCM57_vae_33 decrease with increasing BMI disparity among hosts.

**Fig. S4.**
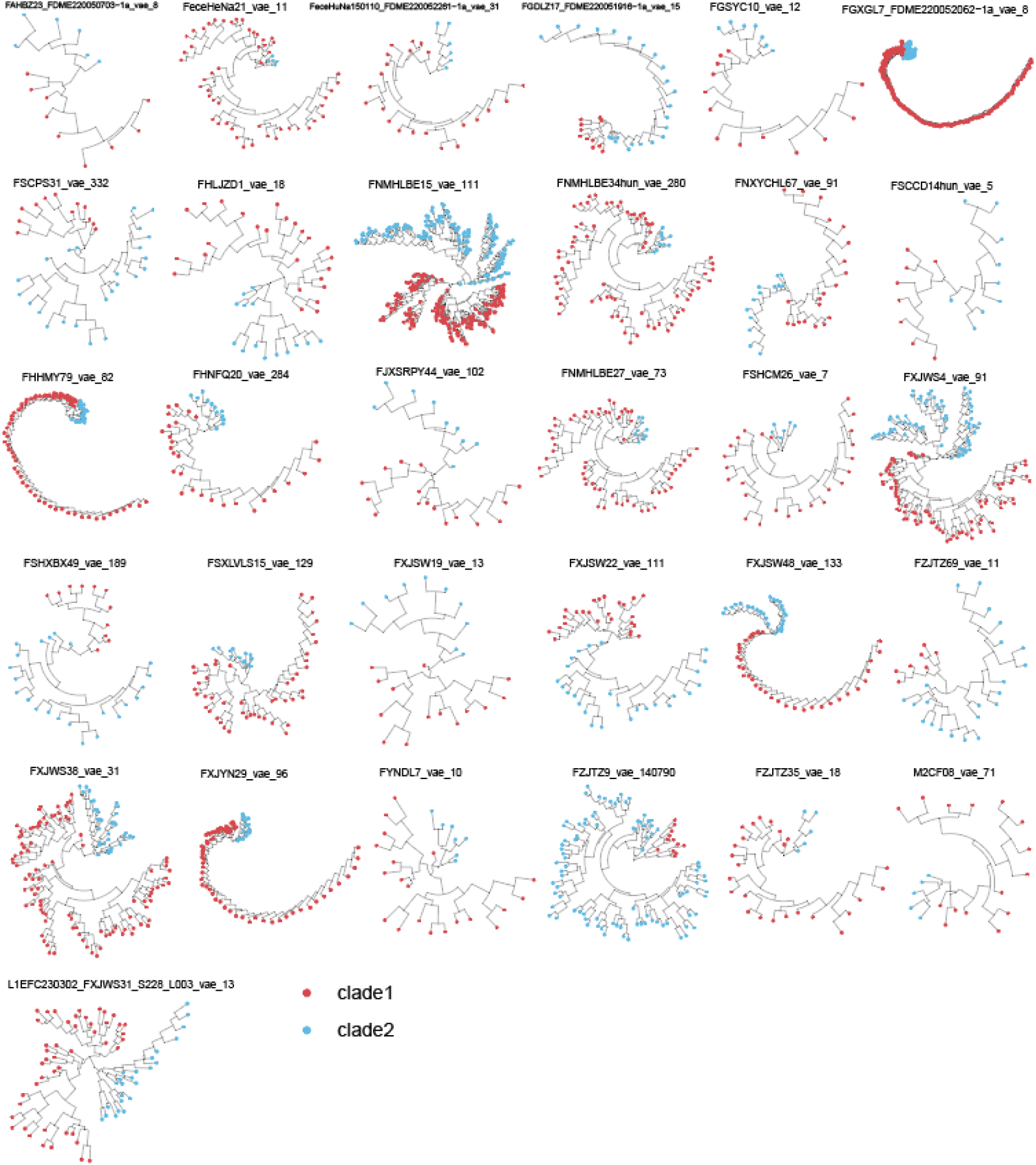
Distinct viral subspecies clades based on their phylogenetic trees. Two Distinct viral subspecies clades detected in 31 VSCs based on their SNP haplotypes. Red and blue represent different subspecies clades.

**Fig. S5.**
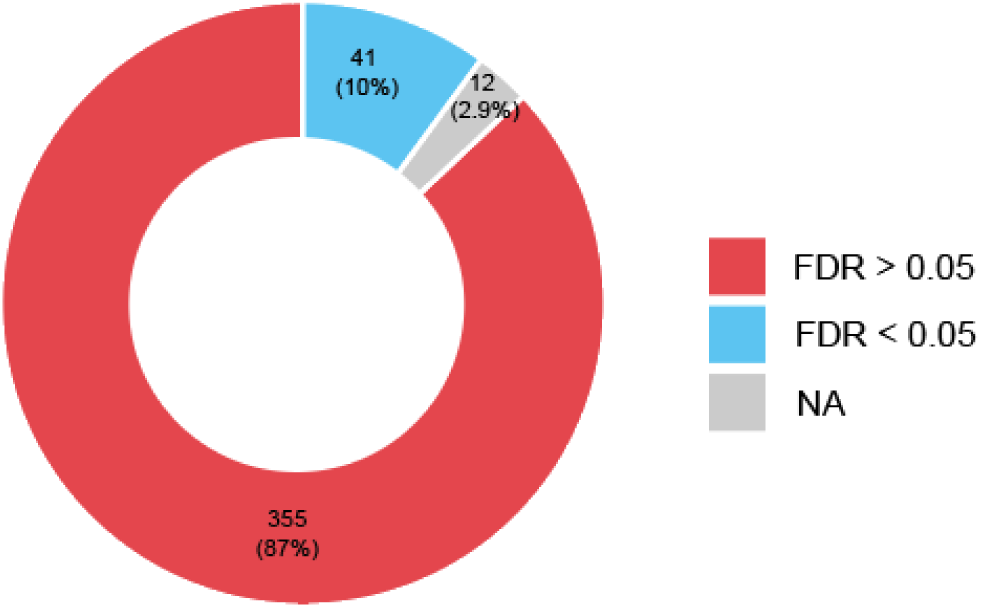
Summary of associations between VSCs genomic dissimilarity and abundance difference. Red indicates proportion of significant associations between VSCs genomic dissimilarity and abundance difference at FDR > 0.05, while blue for FDR < 0.05. NA indicates that the sample size is too small to compute correlations.

**Fig. S6.**
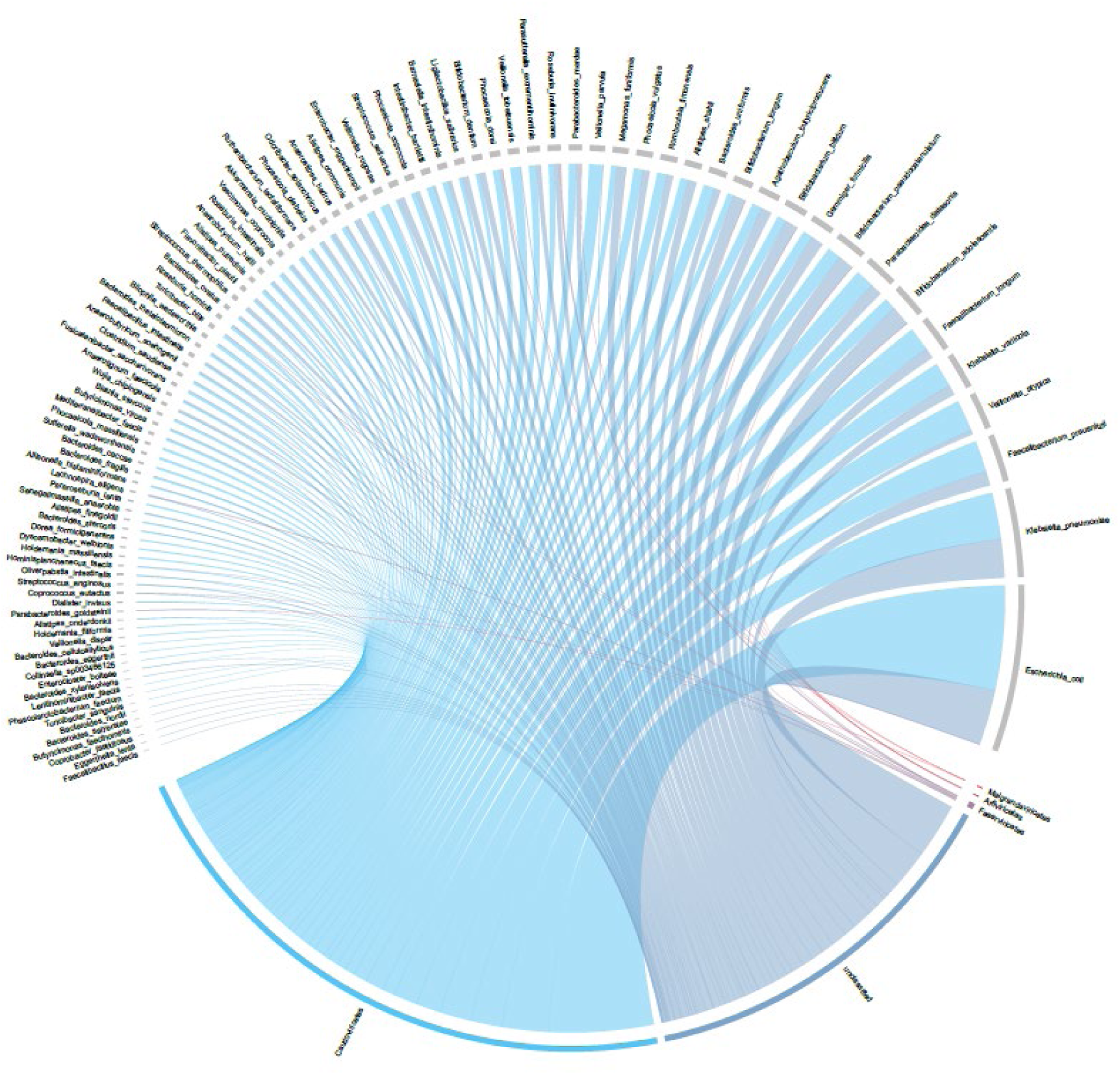
Significant associations between VSCs and bacterial species with CRISPR-spacer based confirmation. In total, 3,497 significant associations between VSCs and bacterial species were characterized, with the most viral associations were observed with *Escherichia coli.*

## Supplementary tables S1 – S18

**Table S1**. The quality of 120,568 medium-to-high quality viral genomes evaluated by CheckV.

**Table S2**. Summary of 64,377 viral genomes from 42,803 VSCs.

**Table S3.** Taxonomy annotation of 42,803 VSCs using ICTV.

**Table S4.** Summary analysis of functional annotation of CDS of 18,311 unclassified VSCs.

**Table S5**. Comparison of genetic distances among different populations of 233 VSCs.

**Table S6**. SNV sites that are significantly different between two populations.

**Table S7.** Matching relationships between viruses and bacteria identified by spacer sequences.

**Table S8**. The intra-group genomic dissimilarity of bacterial hosts corresponding to viruses from different clade.

**Table S9**. SNVs that potentially driving the observed genetic stratification in bacterial hosts.

**Table S10**. The UniRef50 annotation results for driver single nucleotide variants (SNVs).

**Table S11**. Significant associations between viral species genomic and host phenotypic differences.

**Table S12.** Correlations between viral subspecies clades and host phenotypic traits.

**Table S13**. Associations between VSCs genomic dissimilarity and VSC abundance differences.

**Table S14.** Significant associations between viral abundance and phenotypes

**Table S15.** Taxonomic information of 38,436 VSCs involved in the CGVR.

**Table S16.** Significant associations between virus and bacteria.

**Table S17.** Summary of 3,497 vial associations to bacterial species confirmed by CRISPR-spacer imprints.

**Table S18.** The significant results of bacteriome mediate virus impacts on phenotypic traits.

**Correspondence and requests for materials** Further information and requests for resources and reagents should be directed to the lead contact Lianmin Chen.

